# PKA–CIP4 SIGNALING REGULATES CIP4 RELOCATION IN ACTIVATED NATURAL KILLER CELLS

**DOI:** 10.64898/2026.04.22.720117

**Authors:** Alejandro P. Pariani, Victoria Huhn, Leandra Marin, Evangelina Almada, Tomás Rivabella Maknis, Felipe Zecchinati, Rodrigo Vena, Esteban Serra, James R. Goldenring, Cristián Favre, M. Cecilia Larocca

## Abstract

Natural killer (NK) cells are cytotoxic lymphocytes of the innate immune system that eliminate virus-infected and transformed cells through the formation of a specialized immune synapse. Effective target cell killing requires coordinated plasma membrane remodeling and dynamic reorganization of the actin and microtubule cytoskeletons, enabling centrosome polarization and directed secretion of lytic granules. The scaffold protein CIP4 has emerged as an important regulator of cytoskeletal coordination in NK cells, yet how its subcellular localization is controlled during NK cell activation is unknown. CIP4 contains a unique protein kinase A (PKA) phosphorylation site (threonine 225, T225) within its F-BAR domain, a domain that mediates interactions with microtubules and the plasma membrane. We hypothesized that localized PKA signaling controls CIP4 redistribution during immune synapse assembly. To test this hypothesis, we analyzed CIP4 localization and phosphorylation in NK cells engaged with sensitive target cells using biochemical and imaging approaches. We show that NK–target cell interaction enhances PKA activity and promotes phosphorylation of CIP4, coinciding with its delocalization from microtubules and accumulation at the immune synapse. Importantly, this relocalization process requires the PKA-anchoring protein AKAP350, which positions PKA and CIP4 within the same protein complex, thereby facilitating CIP4 phosphorylation. Consistently, pharmacological inhibition of PKA prevented CIP4 delocalization from microtubules and reduced its accumulation at the immune synapse. The non-phosphorylatable CIP4 mutant T225A displayed increased association with microtubules compared with a phosphomimetic mutant, identifying phosphorylation at T225 as a key determinant of CIP4 spatial regulation. Together, these findings identify a signaling mechanism that links compartmentalized PKA activity to the spatial control of CIP4 during immune synapse formation, providing new insight into the molecular mechanisms governing immune synapse maturation.

## 1. INTRODUCTION

Natural killer (NK) cells are cytotoxic lymphocytes of the innate immune system that provide a first line of defense against virus-infected and transformed cells. NK effector function relies on a tightly coordinated cytolytic program driven by dynamic remodeling of the plasma membrane and cytoskeleton (Lagrue et al., 2013). Following interaction with sensitive target cells, NK cells execute directed killing through initiation, effector, and termination phases (Mace et al., 2014). During the initiation phase, NK cells recognize and adhere to target cells, forming an immune synapse (IS) where activating and inhibitory receptor signals are integrated. At this stage, microtubule-dependent transport promotes convergence of lytic granules around the centrosome, the best-characterized microtubule-organizing center (MTOC). When activating signals exceed inhibitory inputs, the effector phase is initiated. This phase involves extensive actin reorganization, driven sequentially by formin- and Arp2/3-mediated polymerization, along with coordinated centrosome repositioning and directional transport of lytic granules toward the IS (Carisey et al., 2018; Mentlik et al., 2010). Subsequent actin clearance and remodeling at the synaptic interface enable granule docking, membrane fusion, and focused secretion of cytotoxic molecules into the target cell. Finally, NK cells disengage and reorient their cytolytic machinery to engage new targets. Although the contributions of membrane remodeling and cytoskeletal dynamics to these processes are well established, the cellular and molecular mechanisms that coordinate cytoskeletal reorganization with membrane remodeling remain incompletely understood (Lee, Wong & Ding, 2021).

CDC42-interacting protein 4 (CIP4) has been identified as an integrator of actin and microtubule reorganization during IS formation in NK cells (Banerjee et al., 2007). CIP4 is a CDC42 effector protein containing an N-terminal F-BAR domain, composed of an FCH (Fes–CIP4 homology) region followed by a coiled-coil segment that mediates dimerization and membrane curvature generation (Aspenström 1997; Shimada et al., 2007). Downstream of the F-BAR domain, CIP4 contains an HR1 motif that binds GTP-bound CDC42 and a C-terminal SH3 domain that recruits key actin-regulatory proteins, including WASp and the formin hDia (Tian et al., 2000; Aspenström et al., 2006). These structural features are consistent with CIP4’s role in processes requiring coordinated regulation of membrane remodeling and actin dynamics, such as endocytosis, which involves membrane invagination, as well as the formation of invadopodia and lamellipodia, which depend on membrane protrusion (Fricke et al., 2009; Pichot et al., 2010; Saengsawang et al., 2012). CIP4 also interacts with microtubules through its FCH domain (Tian et al., 2000), a property that contributes to podosome formation in macrophages (Linder et al., 2000). In NK cells, CIP4 translocates to the centrosome and the IS upon engagement with susceptible targets, where it associates with both the actin and microtubule cytoskeletons. Alterations in CIP4 expression or localization disrupt IS maturation, underscoring the importance of CIP4 positioning during NK activation (Banerjee et al., 2007). However, the mechanisms governing CIP4 localization during NK–target cell interactions remain largely unknown.

Previous work from our group demonstrated that protein kinase A (PKA) selectively phosphorylates CIP4 at residue T225 (Larocca et al., 2004; Tonucci et al., 2019). T225 is located on the convex surface of the F-BAR domain (Tonucci et al., 2019; Shimada et al., 2007). In breast cancer cells, phosphorylation at this site enhances CIP4 recruitment to invadopodia, promotes its interaction with CDC42, and supports invadopodia formation (Tonucci et al., 2019). Whether the PKA–CIP4 signaling axis operates in other cellular contexts has not been explored. Here, we investigate the regulation of CIP4 phosphorylation by PKA during NK cell activation and examine the functional consequences of this modification on CIP4–microtubule interaction and CIP4 relocalization to the IS.

## 2. MATERIALS AND METHODS

### 2.1 Cell lines

The immortalized NK cell line YTS (no unique RRID available to our knowledge; see Gunesch et al., 2019 for genomic characterization) and the KT86 cell line, a derivative of the MHC class I–negative K562 erythroleukemia cell line (RRID: CVCL_0004) stably expressing CD86, were kindly provided by Dr. Jordan Orange (Texas Children’s Hospital, Houston, TX, USA) to Dr. Norberto Zwirner (IBYME-CONICET, Buenos Aires, Argentina) in 2015. Cells were frozen at passages 2–15 after receipt and were used at passages below 25. Cells were maintained in RPMI 1640 medium (Gibco, Thermo Fisher Scientific, Buenos Aires, Argentina) supplemented with 10% fetal bovine serum (FBS; Natocor, Córdoba, Argentina), 1% L-glutamine, 1% nonessential amino acids, and 1% penicillin/streptomycin (all from Gibco, Thermo Fisher Scientific), at 37 °C in a humidified atmosphere containing 5% CO₂.

NK-YTS cells with reduced AKAP350 expression (AKAP350KD) were generated as previously described (Pariani et al., 2023). Briefly, oligonucleotides containing the AKAP350 mRNA target sequence (5′-GCAAGAACTAGAACGAGAA-3′) were annealed and ligated into the AgeI and EcoRI sites of the pLKO.1-puro vector (Addgene). Constructs were verified by sequencing and co-transfected into HEK293FT cells together with the ViraPower lentiviral packaging mix (Invitrogen, Carlsbad, CA, USA). Viral supernatants were collected after 24 h and used to transduce YTS or ex vivo NK cells overnight. Cells were allowed to recover for 24 h and then selected with puromycin (2 µg/mL) for 7 days. AKAP350 silencing was confirmed by Western blot.

C3A cells (HepG2/C3A; RRID: CVCL_1098) were obtained from the American Type Culture Collection (ATCC) by Dr. Pablo J. Schwarzbaum (IQUIFIB, CONICET–Universidad Nacional de Buenos Aires) in 2005. Cells were frozen at passages 4–15 and were used for experiments up to passage 25. Cells were cultured in Dulbecco’s modified Eagle’s medium containing 4.5 g/L glucose, supplemented with 10% FBS and antibiotics. C3A CIP4 T225A and CIP4 T225E stable cell lines were generated and cultured as previously described (Tonucci et al., 2019). All cell lines were tested monthly for mycoplasma contamination by PCR.

### 2.2 Ex vivo NK cell purification

Human samples for NK cell isolation were obtained with informed consent from all donors and used according to protocols approved by the Institutional Review Board of Hospital Provincial del Centenario (Rosario, Argentina). Twelve donors were enrolled. Human NK cells were isolated using the RosetteSep Human NK Cell Enrichment Cocktail (STEMCELL Technologies #15025, Vancouver, Canada) and Ficoll-Paque PLUS (Cytiva, Lobov Scientific, Buenos Aires, Argentina) as previously described (Pariani et al., 2023). Briefly, whole blood was collected using heparin as anticoagulant. To obtain the buffy coat, 20 mL of blood was centrifuged at 900 × g for 15 min. The buffy coat was resuspended in PBS and layered onto a Ficoll gradient, followed by centrifugation at 600 × g for 30 min to isolate PBMCs. PBMCs were washed twice with physiological solution and centrifuged at 400 × g for 15 min. PBMCs were then resuspended in 500 μL of whole blood and incubated for 20 min at room temperature with RosetteSep reagent for negative selection. After incubation, an equal volume of physiological solution containing 2% FBS was added, and the NK-enriched fraction was recovered by centrifugation over a Ficoll gradient (400 × g, 20 min). Purified NK cells were washed twice in physiological solution with 2% FBS (400 × g, 10 min), resuspended in RPMI 1640 medium supplemented as above, and maintained with IL-2 (500 IU/mL; PeproTech, Rocky Hill, NJ, USA) at 37 °C and 5% CO₂ for 24 h to 10 days depending on the experiment.

### 2.3 Target cell–mediated activation of NK cells

KT86 cells were labeled with a vital fluorophore to distinguish them from NK cells. Briefly, KT86 cells were washed with PBS, resuspended at 2 × 10⁶ cells/mL, and incubated with CellTracker™ Deep Red (300 nM; Invitrogen) for 15 min at 37 °C. Labeling was stopped by adding an equal volume of FBS. Cells were washed and resuspended in complete RPMI. NK–target cell conjugates were formed by incubating NK and KT86 cells at a 2:1 ratio in suspension for 15 min at 37 °C, followed by adhesion to poly-L-lysine–coated glass slides (Sigma-Aldrich) for an additional 15 min.

For experiments evaluating NK activation specifically through LFA-1 and CD28 receptors, NK-YTS cells were plated on poly-L-lysine–coated coverslips coated either with nonspecific mouse IgG (resting condition) or with ICAM-1 (5 µg/mL; BioLegend #552906) together with an activating anti-CD28 antibody (5 µg/mL; BD Biosciences #554121) for 30 min at 37 °C, then washed, fixed, stained, and imaged by confocal microscopy as described below.

### 2.4 Assessment of PKA/CIP4 phosphorylation pathway

The activation of the PKA/CIP4 phosphorylation pathway was evaluated in NK-YTS cells exposed to sensitive target cells and in their corresponding non-activated controls. YTS and KT86 cells were mixed at a 6:1 effector:target ratio and centrifuged for 15 s at 100 × g to favor conjugate formation. Cells were incubated for 30 min at 37 °C and 5% CO₂. Non-activated control samples were prepared by incubating YTS and KT86 cells separately for the same period, followed by mixing on ice at a 6:1 ratio and immediate centrifugation. Cells were harvested and lysed in PBS (pH 7.4) containing 1% Triton X-100 and protease and phosphatase inhibitors, then subjected to two freeze–thaw cycles (−80 °C). Activation of PKA signaling was assessed by immunoblot analysis of phosphorylated PKA substrates using an antibody recognizing the RXXS/T(Pi) motif (Cell Signaling #9621, 1:1000), as previously described (Zucchetti et al., 2014). CIP4 phosphorylation was evaluated in CIP4 immunoprecipitates obtained from the same lysates. For immunoprecipitation, cell lysates were centrifuged at 1000 × g for 5 min to remove debris, and supernatants were incubated with a mouse anti-CIP4 antibody (BD Biosciences #612556), as previously described (Larocca et al., 2004). Immunoprecipitates were resuspended in sample buffer (20 mM Tris-HCl pH 8.5, 1% SDS, 400 μM DTT, 10% glycerol), and phosphorylated CIP4 was analyzed as detailed in the Immunoblotting section.

### 2.5 Biochemical assessment of in situ CIP4 association with microtubules

Microtubule-enriched extracts and associated proteins were prepared following a protocol originally described for neuronal cells (Brown et al., 1992). NK-YTS cells were plated on 3-cm dishes coated with either nonspecific mouse IgG (resting condition) or ICAM-1 (5 μg/mL) together with an activating anti-CD28 antibody (5 μg/mL) and incubated overnight. Cells were washed with PBS and then with PHEM buffer (60 mM PIPES, 25 mM HEPES, 10 mM EGTA, 2 mM MgCl₂, pH 6.9). To stabilize polymerized microtubules and extract soluble or non–microtubule-bound proteins, cells were permeabilized for 5 min with PHEM containing 1% Triton X-100, 10 μM taxol, and 0.1% DMSO in the presence of protease inhibitors. The extraction buffer containing soluble proteins was collected. Cells were then washed with detergent-free PHEM and scraped into 100 μL of sample buffer. Both fractions (soluble proteins and microtubule-associated proteins) were analyzed by Western blot. The percentage of CIP4 associated with microtubules was calculated as the amount of CIP4 in the microtubule-enriched fraction relative to the total CIP4 present in both the microtubule-enriched and soluble fractions. To account for potential differences in total microtubule recovery resulting from changes in microtubule stability during NK-cell activation, CIP4 levels in the microtubule-enriched fraction were additionally normalized to α-tubulin density for each experimental condition.

### 2.6. In silico assessment of CIP4 T225 and CIP4 T225-phosphorylated dimer properties

To assess the impact of CIP4 T225 phosphorylation on CIP4 dimerization, structural predictions of the CIP4 F-BAR domain were generated using AlphaFold Server (Jumper et al., 2021). Models were computed for both non-phosphorylated and phosphorylated forms of the F-BAR domain, with phosphorylation at residue T225 introduced in silico. For each condition, dimeric models were generated and evaluated using the inter-chain predicted TM (ipTM) score. Higher ipTM values correlate with increased confidence in global structure, and comparison of ipTM scores across models have been used to assess relative differences in predicted interface quality (O’Reilly et al., 2023). In parallel, pLDDT estimates AlphaFold 3’s confidence for atoms position in the structure prediction and the predicted aligned error (PAE) graphic was used as a measure for confidence in the relative positions of two monomers within the predicted structure of the dimmers. The PAE matrix length corresponds to AlphaFold-Multimer internal indexing and may exceed the number of biological residues due to the inclusion of chain separators.

### 2.7 Immunoblotting

Protein concentration in cell lysates was determined according to Lowry et al. (1951). Equal amounts of protein (cell lysates) or equal sample volumes (immunoprecipitates in Figure 1; microtubule-enriched and soluble fractions in Figures 2 and 3) were heated for 10 min at 90°C in sample buffer and resolved on SDS–10% gels (Figures 1–3) or on SDS 4–10% gradient gels (Figure 5). Proteins were transferred to polyvinylidene difluoride (PVDF) membranes (PerkinElmer Life Sciences; Figures 1–3) or nitrocellulose membranes (Amersham Pharmacia Biotech; Figure 5). Membranes were blocked with 5% non-fat dry milk in PBS containing 0.3% Tween-20 (PBS-Tween). PVDF membranes were probed with rabbit anti-RXXS/T(Pi) (Cell Signaling, 1:1000), mouse anti-CIP4 (BD Biosciences #612556, 1:500), or mouse anti-α-tubulin (Sigma T5168, 1:5000). Nitrocellulose membranes for AKAP350 detection were probed with the mouse monoclonal anti-AKAP350 (14G2) (Schmidt et al., 1999) or anti-α-tubulin. After washing, membranes were incubated with the appropriate horseradish-peroxidase–conjugated secondary antibodies, and signals were detected by enhanced chemiluminescence (Pierce, Thermo Scientific). Images were acquired using an Amersham ImageQuant 500 system (Cytiva). Band intensities were quantified by densitometry using NIH ImageJ.

**Figure 1.**
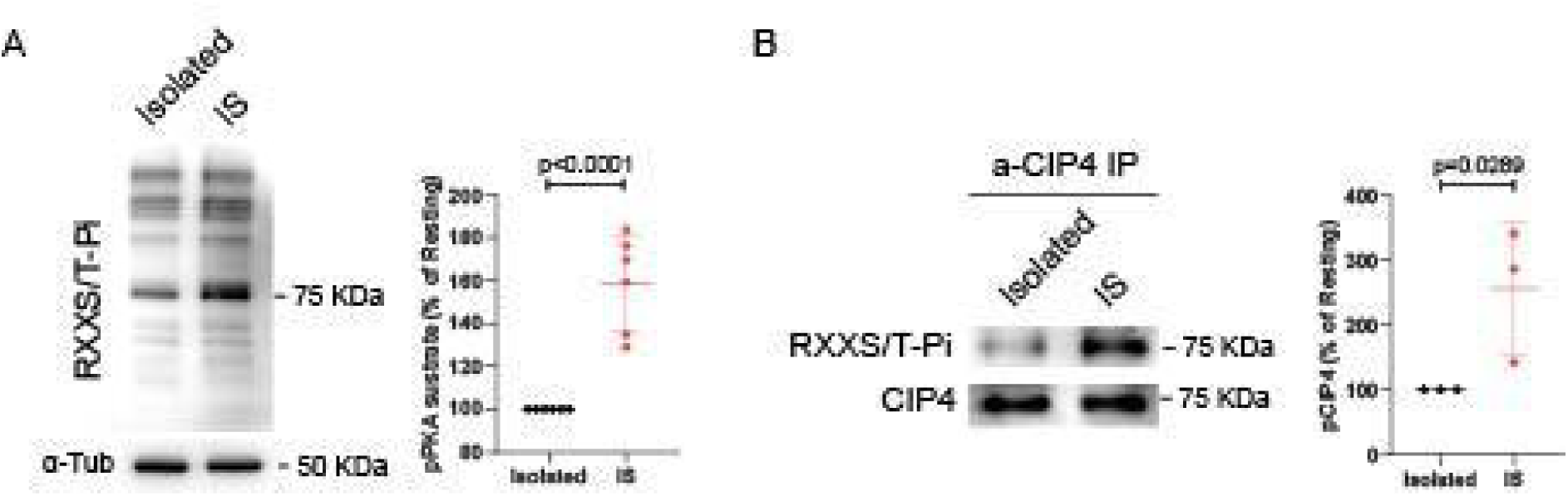
NK–target cell interaction induces PKA activation and CIP4 phosphorylation. YTS NK cells were incubated in absence (Isolated) or in presence (IS) of KT86 target cells, for 30 min as described in Materials and Methods. A) Whole-cell lysates were analyzed by immunoblotting using an antibody recognizing phosphorylated PKA substrates (RXXS/T(Pi) motif). Representative immunoblots for RXXS/T(Pi) and α-tubulin, used as a loading control, are shown. Dot plots depict densitometric quantification of PKA substrate phosphorylation normalized to α-tubulin from six independent experiments. B) CIP4 immunoprecipitates were prepared, and CIP4 phosphorylation was evaluated by immunoblotting using the RXXS/T(Pi) antibody. Representative immunoblots for phosphorylated CIP4 and total CIP4 are shown. Dot plots represent densitometric quantification of CIP4 phosphorylation normalized to total CIP4 from three independent experiments. Data are presented as mean ± SEM.

**Figure 2.**
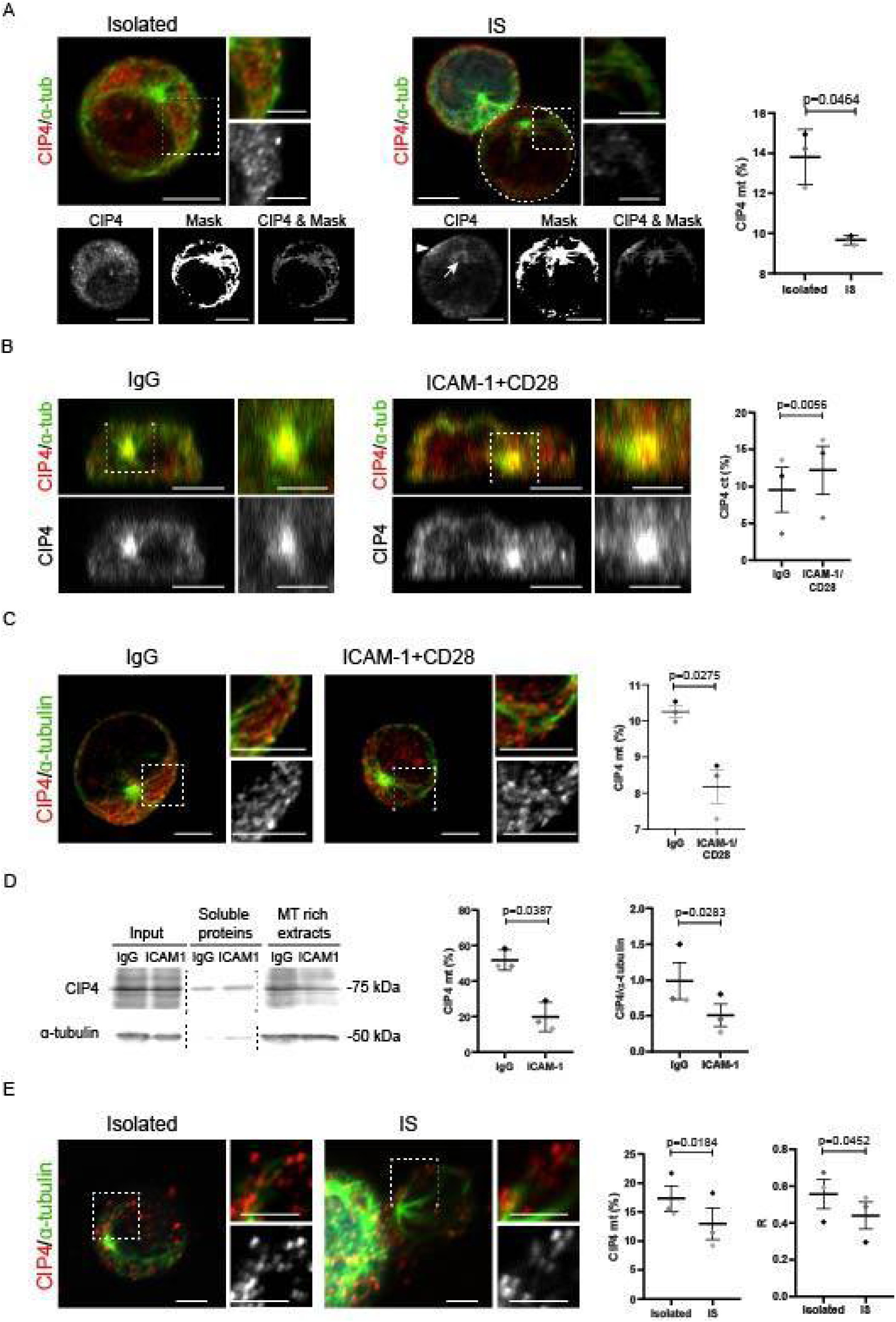
NK–target cell interaction induces CIP4 delocalization from microtubules. A and E) KT86 cells were labeled with CellTracker™ Deep Red to allow their discrimination from NK cells, and NK–target cell conjugates were formed by incubating NK and KT86 cells at a 2:1 ratio for 30 min, as described in the Materials and methods section. Cells were then fixed, stained, and analyzed by confocal microscopy. Images were obtained as described in Materials and methods. A) Images show merged staining for CIP4 (red) and α-tubulin (green) in isolated NK-YTS cells or in conjugates labeled with CellTracker (cyan) (upper panels); and the CIP4 channel, the mask generated using the α-tubulin channel, and the result of applying the ImageJ *Image Calculator/AND* tool to these images to estimate the fraction of CIP4 associated with microtubules (lower panels). Insets show magnifications of the boxed areas in the merged images and in the CIP4 channel (grayscale). The dot plot shows quantification from three independent experiments of the percentage of CIP4 associated with microtubules in isolated NK-YTS cells and in NK-YTS conjugated to target cells. B–C) NK-YTS cells were plated on poly-L-lysine–coated coverslips coated either with nonspecific mouse IgG (resting condition) or with ICAM-1 together with an activating anti-CD28 antibody (activated condition) and incubated for 30 min at 37 °C. Cells were then washed, fixed, stained, and imaged by confocal microscopy. Images show merged staining for CIP4 (red) and α-tubulin (green) in cells incubated under resting or activated conditions (upper panels), and the corresponding CIP4 channel in grayscale (lower panels) for orthogonal x,z views of z-stack reslices (B). Insets show magnifications of the boxed areas. Dot plots show quantification from three independent experiments of the percentage of CIP4 associated with centrosomes (B), or with microtubules (C), in resting and activated NK-YTS cells. D) NK-YTS cells were plated on 3-cm dishes coated either with nonspecific mouse IgG (resting condition) or with ICAM-1 (5 μg/mL) together with an activating anti-CD28 antibody (5 μg/mL) (activated condition) and incubated for 30 min. Microtubule-enriched fractions and associated proteins were prepared as detailed in Materials and Methods, and the presence of CIP4 in the microtubule-enriched fraction and in the remaining lysate was analyzed by immunoblotting. Images show representative immunoblots for CIP4 and α-tubulin, the latter used to verify microtubule enrichment in each fraction. The dashed line indicates lanes from the blot that were omitted in the figure for clarity. The dot plots show quantification of the percentage of CIP4 associated with microtubules, and of CIP4 levels in the microtubule-enriched fraction normalized to α-tubulin density from three independent experiments. E) Images show merged staining for CIP4 (red) and α-tubulin (green) in isolated ex vivo NK (exNK) cells or in conjugates of exNK cells with target cells (cyan). Insets show magnifications of the boxed areas in the merged images and in the CIP4 channel (grayscale). The dot plot shows quantification from three independent experiments of the percentage of CIP4 associated with microtubules in isolated exNK cells and in exNK cells conjugated to target cells. At least eight cells were analyzed per experiment (A-C and E). Data are presented as mean ± SEM. Each dot in the dot plots represents an independent experiment, shown in a different color to indicate paired data. Scale bars: 5 µm (A-main panels, C) and 2.5 µm (A-insets, B, E).

**Figure 3.**
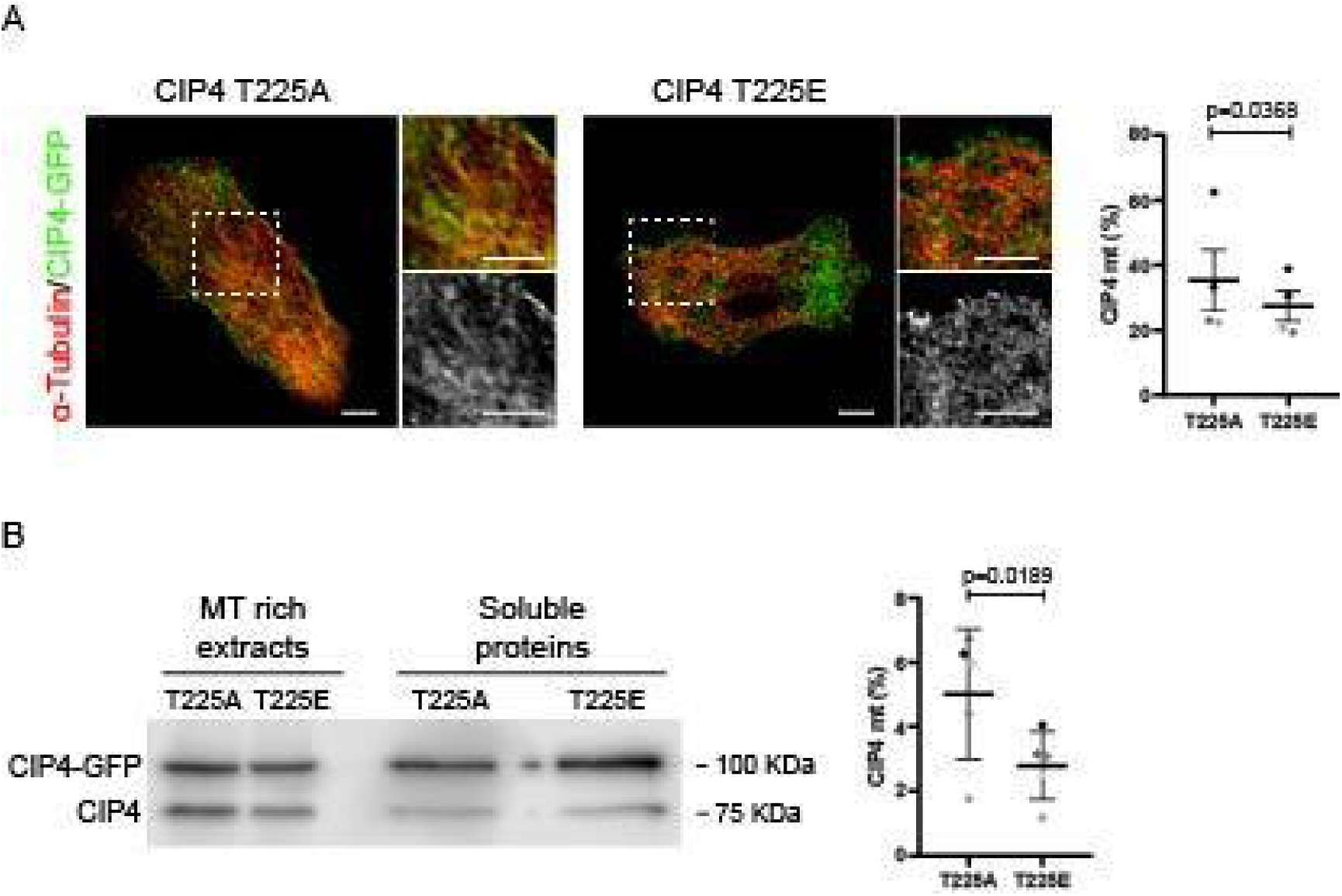
CIP4T225E shows reduced association with microtubules compared with CIP4T225A. C3A cells stably expressing CIP4 T225A or CIP4 T225E as GFP fusion proteins were fixed, stained for α-tubulin, and analyzed by confocal microscopy. A) Images show merged staining of α-tubulin (red) and GFP fluorescence indicating the expression of each CIP4 mutant (green). Insets show higher-magnification views of the merged staining and the GFP channel (grayscale) from the boxed areas. The dot plot shows quantification from four independent experiments of the percentage of CIP4 fusion protein associated with microtubules for each mutant, assessed as shown in Figure 2A. At least eight cells were analyzed per experiment. Scale bars, 5 µm. B) Microtubule-enriched fractions and associated proteins were prepared, and levels of CIP4 proteins in each fraction were analyzed by inmunoblot. The image shows a representative immunoblot for CIP4, including bands corresponding to endogenous CIP4 (75 kDa) and the fusion protein (100 kDa, CIP4-GFP). The dot plot shows quantification of the percentage of CIP4T225A and CIP4T225E associated with microtubules from four independent experiments. Each dot in the dot plots represents an independent experiment, shown in a different color to indicate paired data. Data are presented as mean ± SEM.

### 2.8 Immunofluorescence

Cells from each experimental condition were fixed, permeabilized, and blocked prior to incubation with the corresponding primary antibodies for 2 h. After washing, coverslips were incubated for 1 h with secondary antibodies conjugated to Alexa 488, Alexa 560, or Alexa 633 (Molecular Probes A34055, 1:200, Thermo Fisher, CABA, Argentina) and with 4′,6-diamidino-2-phenylindole (DAPI; Thermo Scientific, Thermo Fisher, CABA, Argentina). Samples were mounted with ProLong (Thermo Fisher, CABA, Argentina). Images were acquired using an LSM880 confocal system coupled to an Observer Z1 inverted microscope. To optimize visualization and quantification of CIP4 association with microtubules, the ImageJ “stack projection/average” tool was applied to the α-tubulin and CIP4 channels to reduce non-specific fluorescence. For image acquisition, the x–y plane exhibiting maximal α-tubulin signal was selected, and a time-series of 100 frames was obtained at 2.53-s intervals. Average projections were then generated for both α-tubulin and CIP4 channels. For analysis of CIP4 localization at the IS, serial optical sections (0.35 µm) were collected along the z-axis. Z-stacks were reconstructed, and for each NK–KT86 cell pair, the x–y plane providing optimal visualization of the IS was selected for quantification. For analysis of centrosomal CIP4 in ICAM-1/anti-CD28–activated cells, serial optical sections (0.35 µm) were similarly obtained, and x–z reslices were generated using the ImageJ “reslice” tool. For the preparation of the final figures, brightness and contrast adjustments were applied uniformly to the entire image to improve fluorescence visualization.

### 2.9 Image Analysis

CIP4 localization on microtubules. Masks for α-tubulin and CIP4 were generated from the average projections of each channel. For α-tubulin, additional processing with the ImageJ “sharpen” tool was applied prior to binarization to better delineate individual microtubules.

CIP4 levels associated with microtubules and total cellular CIP4 were quantified using the ImageJ Image Calculator/AND logical function. The percentage of CIP4 associated with microtubules was calculated for each cell. Alternatively, in ex vivo NK cells, which volume is about 3 times lower that NK-YTS cells, and the precision of the mask for selecting microtubule ROIs is reduced, we additionally analyzed CIP4 colocalization with microtubules by using the ImageJ colocalization test, and comparing the Pearsońs correlation coefficient for both isolated *ex vivo* NK cells, and NK cells that were interacting with sensitive target cells.

CIP4 localization at the IS. The IS was defined as the cell–cell contact region identified in differential interference contrast (DIC) images. Using the same DIC images, the perimeter of the YTS cell forming the IS was manually outlined (Figure 4). Total fluorescence intensity was measured at the IS and in the whole cell, and the percentage of CIP4 enrichment at the IS was calculated.

**Figure 4.**
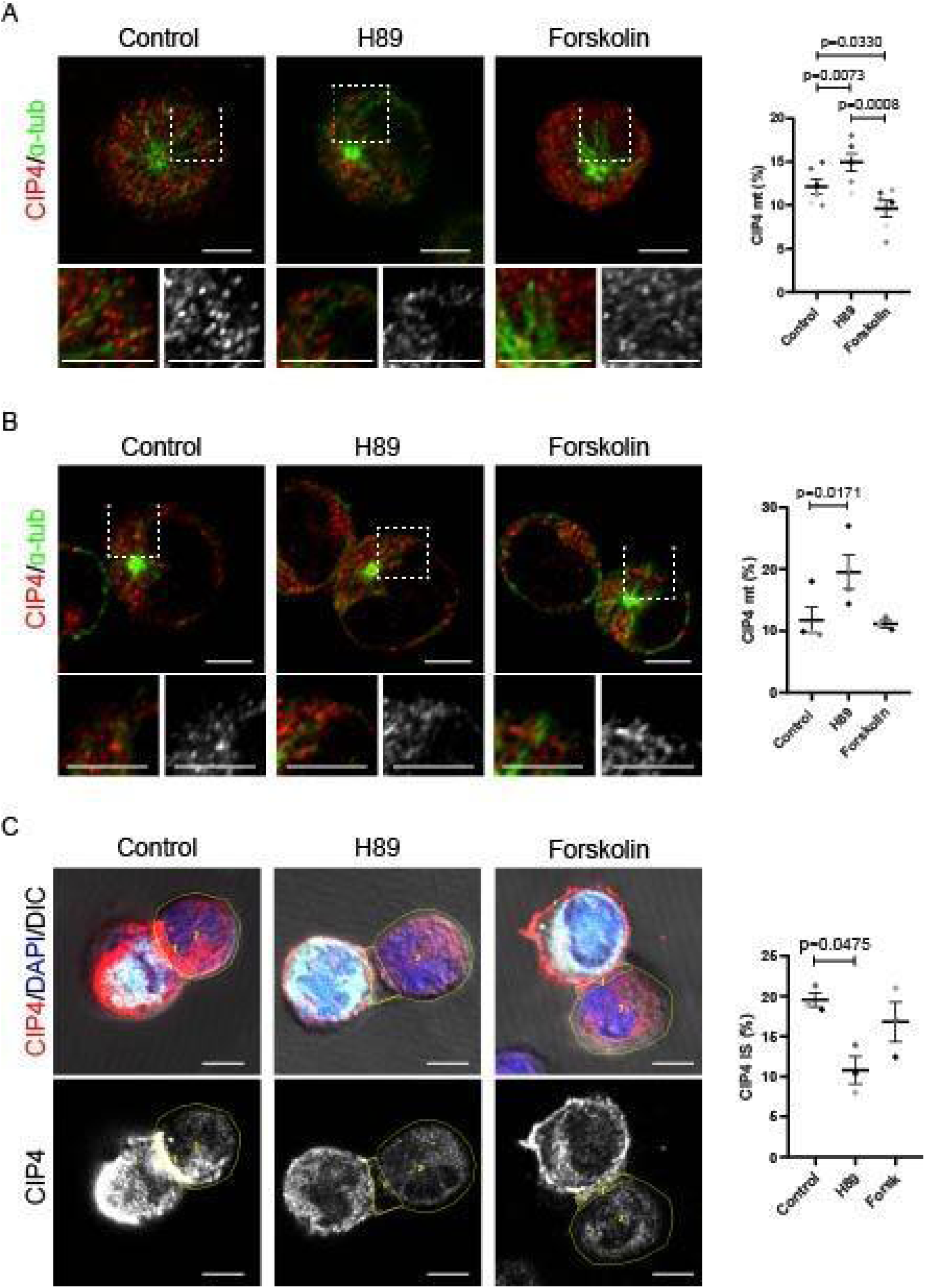
PKA signaling is required for CIP4 reorganization during NK cell activation. NK-YTS and KT-86 cells were incubated for 30 min as described in Figure 2A under control conditions or in the presence of H89 (Santa Cruz, 1 µM) or forskolin (Santa Cruz, 5 µM). Cells were then fixed, stained, and analyzed by confocal microscopy. A, B) Images show merged staining of CIP4 (red) and α-tubulin (green) in isolated NK cells (A) or in conjugates with target cells (B). Insets show magnifications of the boxed areas in the merged images and in the CIP4 channel (grayscale). Dot plots show quantification from four (A) or three (B) independent experiments of the percentage of CIP4 associated with microtubules in isolated NK-YTS cells (A) and in NK cells conjugated to target cells (B). C) Images show merged DIC (grayscale), CIP4 (red), nuclear DAPI (blue) and cell tracker staining of KT86 cells (cyan) channels at the z level where the immunological synapse (IS) was best visualized (upper panels), and the CIP4 channel in grayscale (lower panels). ROIs corresponding to the IS and to the NK-YTS cell contour are outlined in yellow. The dot plot shows quantification of the percentage of CIP4 present at the IS from three independent experiments. At least 8 cells were analyzed per experiment (A–C). Data are presented as mean ± SEM. Each dot in the dot plots represents an independent experiment, shown in a different color to indicate paired data. Scale bars, 5µm.

CIP4 localization at the centrosome. The region of interest (ROI) corresponding to the centrosome was delimited at the α-tubulin channel using the x–z reslices of the z-stacks in the plane where the centrosome was most clearly visualized, and the ROI corresponding to total CIP4 cell staining was delimited at the same plane using the CIP4 channel. Fluorescence intensity in the CIP4 channel was measured within each ROI, and the percentage of centrosomal CIP4 was calculated for each cell.

### 2.10 Statistical analysis

Data are expressed as mean ± SEM. Paired Student’s t-test or nonparametric Mann–Whitney test were used for comparison between groups when necessary. For multiple comparisons, 1-way ANOVA followed by Tukey’s multiple comparisons test was used. P < 0.05 was considered statistically significant.

## 3. RESULTS

### 3.1. NK cell activation induces CIP4 phosphorylation by PKA

Residue T225 of CIP4 is located within the F-BAR domain, which mediates CIP4 dimerization and membrane interaction (Shimada et al., 2007) as well as microtubule binding (Tian et al., 2000). To assess whether CIP4 phosphorylation is regulated during NK cell activation, we examined activation of the PKA–CIP4 phosphorylation pathway upon NK–target cell engagement. Consistent with previous reports showing that interaction with sensitive target cells increases intracellular cAMP levels in NK cells (Whalen et al., 1998), immunoblot analysis of NK-YTS cell lysates revealed that co-culture with sensitive target cells induced a marked increase in the immunoreactivity of several proteins detected with an antibody recognizing phosphorylated PKA substrates (RXXS/T-Pi motif; Figure 1A), consistent with the activation of a PKA signaling pathway during NK–target cell engagement. To directly assess CIP4 phosphorylation under these conditions, endogenous CIP4 was immunoprecipitated and its phosphorylation was evaluated using the phospho-PKA substrate antibody. NK-YTS cells engaged with sensitive target cells exhibited a significant increase in CIP4 phosphorylation (+150%) compared with isolated cells. Consistent with our previous findings showing that mutation of T225 abolishes PKA-mediated phosphorylation of CIP4 (Tonucci et al., 2019), T225 represents the sole PKA phosphorylation site in CIP4. Furthermore, to our knowledge CIP4 T225 is the only residue detected by the phospho-PKA substrate antibody. Therefore, those results indicate that NK interaction with sensitive target cells induces CIP4 T225 phosphorylation by PKA.

### 3.2. NK cell activation induces a decrease in CIP4 co-localization with microtubules

Banerjee et al. (2007) reported that, during the interaction with sensitive target cells, NK cells showed increased CIP4 colocalization with α-tubulin at the centrosome, next to the IS area. In order to complete the characterization of the dynamics of CIP4 association with microtubules in NK cells, we first characterized CIP4 localization at microtubules in both resting and activated NK cells. Immunofluorescence analyses showed that approximately 14% of total CIP4 colocalized with microtubules in isolated NK-YTS cells and that this localization was significantly reduced upon engagement with sensitive target cells (Figure 2A).

To further characterize CIP4 dynamics under defined stimulatory conditions, we examined its colocalization with microtubules in NK-YTS cells incubated either with non-activating mouse IgG (resting cells) or with the LFA-1 ligand ICAM-1 together with an activating anti-CD28 antibody. Centrosome polarization toward the activating surface was confirmed in x,z reslices generated from z-stacks of each field (Figure 2B). Consistent with the findings of Banerjee et al. (2007) in target cell–stimulated NK cells, analysis of these reslices revealed that LFA-1/CD28 activation also increased CIP4 localization at the centrosome (Figure 2B). Immunofluorescence analysis showed that, as observed during interactions with sensitive target cells, LFA-1/CD28-mediated stimulation induced a marked reduction in CIP4 association with microtubules (Figure 2C). To further validate CIP4 delocalization from microtubules upon NK-cell activation, we prepared microtubule-enriched fractions from NK-YTS cells subjected to the same stimulatory conditions and quantified CIP4 levels in both the microtubule-enriched and soluble fractions by western blot. In agreement with the immunofluorescence results, activation induced a significant decrease in the amount of CIP4 associated with microtubules (Figure 2D).

Finally, we extended the analysis of the dynamics of CIP4 association to microtubules to ex vivo human NK cells. Cultures derived from four healthy donors displayed a fraction of CIP4 associated with microtubules comparable to that observed in NK-YTS cells, whereas interaction with sensitive target cells induced CIP4 delocalization from microtubules in these primary cells, further supporting the physiological relevance of our observations (Figure 2E). Similar results were obtained when CIP4 colocalization with microtubules was analyzed by ImageJ colocalization test, which indicated that the Pearson’s correlation coefficient between CIP4 and α-tubulin stainings was significantly reduced in ex vivo NK cells that were interacting with sensitive target cells (Figure 2E).

### 3.3. CIP4 T225 phosphomimetic mutant exhibits lower microtubule co-localization than the non-phosphorylatable mutant

Given that we previously characterized a bonafide, single PKA phosphorylation site in CIP4 (Tonucci et al., 2019), we then turned our attention to the possibility that this phosphorylation had an impact on CIP4 association with microtubules. To assess this, we examined the *in situ* behavior of CIP4 mutants in which T225 was replaced either with a non-phosphorylatable alanine (CIP4 T225A) or with a phosphomimetic glutamic acid (CIP4 T225E). Analysis of GFP fusion proteins stably expressed in C3A cells revealed that CIP4 T225E exhibited a significantly lower degree of colocalization with microtubules compared with CIP4 T225A (Figure 3A). Western blot analysis of microtubule-enriched fractions supported these observations: approximately 5% of CIP4 T225A–GFP was recovered in the microtubule-associated fraction, whereas only ∼3% of CIP4 T225E–GFP was detected (Figure 3B). Together, these results indicated that phosphorylation at T225 promotes CIP4 delocalization from microtubules.

CIP4 T225 is located on the convex surface of the F-BAR domain, which mediates dimer formation. Dimeric models generated using the AlphaFold server did not reveal major conformational rearrangements upon introduction of phosphorylation at T225. However, phosphorylation was associated with a marked increase in the ipTM score, from 0.40 in non-phosphorylated CIP4 T225 F-BAR dimers to 0.77 in phosphorylated CIP4 T225 F-BAR dimers, for the best model. pLDDT per-atom, which estimate AlphaFold 3’s confidence in a structure prediction, were also higher for the CIP4 T225 F-BAR dimer than for the non-phosphorylated one (Supplementary Figure 1 and 2). Consistently, the PAE graphic shows homogeneously reduced values all along the phosphorylated models, indicating more confidence for the relative predicted position between chains compared with the non-phosphorylated dimers. These parameters were similar for the five best models obtained in each simulation. Together, these observations indicate that phosphorylation at CIP4 T225 is associated with increased confidence in the predicted dimer interface and a more conformationally constrained dimeric arrangement.

### 3.5. PKA–AKAP350-CIP4 signaling is required for CIP4 reorganization during NK cell interaction with sensitive target cells

Having provided evidence supporting the hypothesis that phosphorylation at T225 facilitates CIP4 delocalization from microtubules in cultured cells, we next examined whether manipulating PKA activity would similarly regulate CIP4 redistribution during NK cell engagement to sensitive target cells. To address this, we first analyzed the effects of the PKA inhibitor H89 and the PKA activator forskolin on CIP4 localization. Treatment with H89 increased, whereas forskolin treatment reduced CIP4 association with microtubules in isolated NK-YTS cells (Figure 4A). In NK-YTS cells forming an IS with sensitive target cells, H89 treatment also increased CIP4 association with microtubules, whereas forskolin did not produce a noticeable effect (Figure 4B). Additionally, the comparison of CIP4 association to microtubules in activated vs. isolated NK cells indicated that PKA inhibition prevented CIP4 delocalization from microtubules (Supplementary Figure 3A). Concomitantly, analysis of CIP4 distribution at the IS revealed that, in the same conditions where H89 increased CIP4 localization at microtubules, H89 also diminished CIP4 accumulation at the IS (Figure 4C). On the other hand, forskolin did not significantly alter CIP4 relocation to the IS (Figure 4C). Altogether, these observations indicate that, during NK cell interaction with sensitive target cells, PKA activity is required for CIP4 delocalization from microtubules and its subsequent accumulation at the IS.

CIP4 T225 phosphorylation is facilitated by the A-kinase anchoring protein 350 (AKAP350; also known as AKAP450 or CG-NAP), which recruits both PKA and CIP4 into the same multiprotein complex (Schmidt et al., 1999; Larocca et al., 2004; Tonucci et al., 2019). To assess the contribution of the assembly of this complex to CIP4 redistribution, we generated NK-YTS cells with reduced AKAP350 expression using a lentiviral shRNA system. AKAP350 knockdown efficiency was confirmed by western blot (Figure 5A). Analysis of CIP4 colocalization with microtubules in IS-forming cells showed that AKAP350 KD cells exhibited a higher fraction of CIP4 associated with microtubules (Figure 5B). Moreover, similar to the phenotype observed with H89 treatment, AKAP350 deficiency resulted in the prevention of CIP4 delocalization from microtubules (supplementary Figure 3B) and reduced CIP4 accumulation at the IS (Figure 5C).

**Figure 5.**
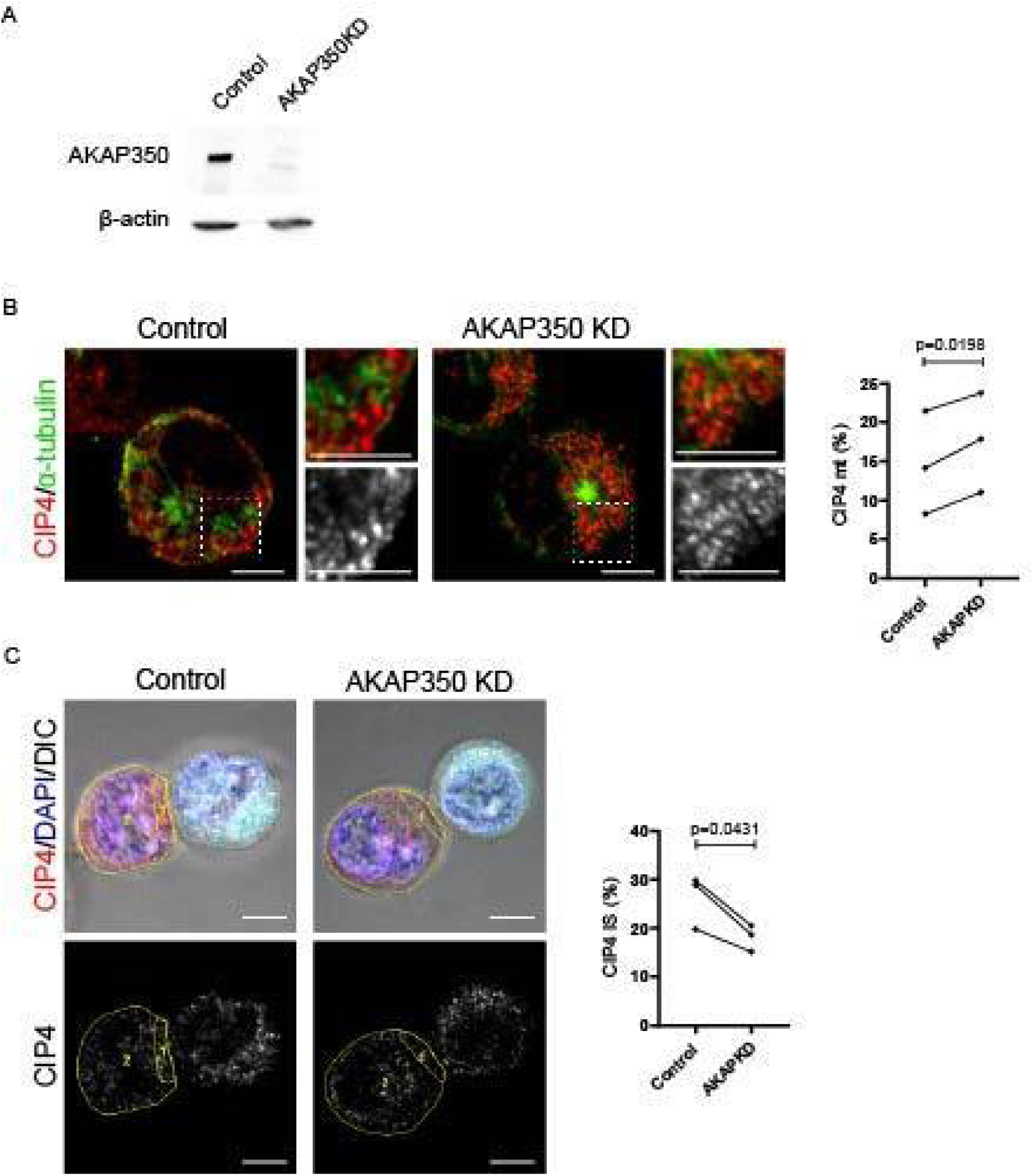
AKAP350 is required for CIP4 reorganization during NK cell interaction with sensitive target cells. NK-YTS cells with decreased AKAP350 expression were generated using a lentiviral system for shRNA expression, as described in the Materials and Methods section. A) Western blot analysis of AKAP350 expression in control and AKAP350KD NK-YTS cell lines. α-tubulin was used as a loading control. B–C) Control and AKAP350KD NK-YTS cells were incubated for 30 min with KT86 cells as described in Figure 2A. B) Images show merged staining of CIP4 (red) and α-tubulin (green) in YTS–KT86 cell conjugates. Insets show magnifications of the boxed areas in the merged images and in the CIP4 channel (grayscale). Dot plots show quantification, from three independent experiments, of the percentage of CIP4 associated with microtubules in NK-YTS cells. Dots from the same experiment are connected by lines to facilitate visualization of paired data. C) Images show merged DIC (grayscale), CIP4 (red), nuclear DAPI (blue), and CellTracker staining of KT86 cells (cyan) at the z-plane where the immunological synapse (IS) was best visualized (upper panels); and the CIP4 channel in grayscale (lower panels). ROIs corresponding to the IS and to the NK-YTS cell contour are outlined in yellow. The dot plot shows quantification of the percentage of CIP4 present at the IS from three independent experiments. Dots from the same experiment are connected by lines to facilitate visualization of paired data. At least eight cells were analyzed per experiment (B, C). Scale bars, 5 µm.

Taken together, these findings support a model in which, during NK cell activation, AKAP350-dependent PKA phosphorylation of CIP4 is required for its release from microtubules and its relocalization to the IS.

## DISCUSSION

During the establishment of the NK-IS, plasma membrane remodeling, localized actin polymerization and centrosome reorientation toward the target cell shape the architecture required for the polarized secretion of the lytic granules towards the target cell. Within this highly orchestrated framework, because of its capacity to interact with the plasma membrane and to bind microtubules and core actin remodeling regulators that have a central role in NK cell cytotoxic response such as CDC42 (Aspenström, 1997, Sinai et al., 2010), WASP (Tian et al., 2000, Orange et al., 2002), and mDIA (*Aspenström, 2007,* Butler et al., 2009), CIP4 stands out as an essential scaffold protein that bridge microtubules and actin regulatory machinery to ensure proper cytoskeletal coordination at the IS to drive the activation of NK cytolytic response (Banerjee et al., 2007). Our findings now reveal that activation of NK cells enhances a PKA/CIP4 phosphorylation pathway that induces CIP4 delocalization from microtubules and recruitment to the IS. Together with the analysis of CIP4 T225 non phosphorylable and phosphomimetic mutants’ association to microtubules, those results support a model in which T225 phosphorylation promotes CIP4 relocalization during the establishment of the IS, thereby facilitating its function in coordinating cytoskeletal organization at the IS during NK cell activation.

Banerjee et al. (2007) showed that CIP4 accumulates at the centrosome and at the IS in NK cells that are compromised in a cytotoxic response and demonstrated that depletion of CIP4 by siRNA impairs IS maturation and NK cytotoxicity, without affecting F-actin accumulation at the IS, thereby positioning CIP4 as a late regulator of IS maturation. Notably, either overexpression of full-length CIP4 or expression of constructs lacking CIP4 microtubule binding domain resulted in aberrant redistribution of CIP4 at the actin-rich cortex, also leading to impairment of IS maturation and reduced cytotoxicity, thus highlighting the relevance of the fine control of CIP4 subcellular localization for NK cytotoxic response. In this context, our results provide a mechanistic framework for understanding how this relocalization is regulated, by identifying a signaling pathway that actively controls the subcellular localization of CIP4 during NK cell activation.

cAMP-PKA pathway has pleiotropic effects on immune cells, and may inhibit NK cytotoxic response (Zhang et al., 2024). Our results add an additional layer of complexity to the classical view of cAMP–PKA signaling as primarily inhibitory for NK-cell effector functions. Two major types of PKA, PKA type I and PKA type II, display different biochemical properties due to differences in their R subunits (RI and RII, respectively). Whereas the activation of PKA type I, which is the major PKA type expressed in NK cells (Torgersen et al 1997), has consistently been linked to decreased cytotoxicity in NK cells exposed to immunosuppressive agonists such as PGE or adenosine (Goto et al., 1983, Torgersen et al., 1997, Lokshin et al., 2006), NK interaction with sensitive target cells have also been shown to induce elevations in cAMP that are required for optimal cytotoxic responses (Whalen et al., 1998, Bariagaber et al., 2003). In this context, we found that NK-cell interaction with sensitive targets triggers an increase in the phosphorylation of PKA substrates, and specifically enhances CIP4 phosphorylation. Furthermore, we found that inhibition of the PKA-CIP4 pathway in NK cells in contact with sensitive target cells through pharmacological blockade with H89 resulted in increased CIP4 association to microtubules and diminished CIP4 accumulation at the IS. We previously showed that, by interacting with PKA-RII subunit, AKAP350 scaffolds PKA type II within a multiprotein complex that promotes CIP4 phosphorylation (Schmidt et al 1999, Larocca et al 2004, Tonucci et al 2019). Given that signal specificity within the cAMP–PKA pathway largely depends on the spatial segregation imposed by AKAPs (Omar and Scott JD, 2020), we hypothesize that localized activation of PKA type II drives cytoskeletal remodeling events necessary for IS assembly. Consistent with this idea, here, we demonstrate that reduced AKAP350 expression, prevents CIP4 delocalization from microtubules and its accumulation at the IS, thus impairing its reorganization during cytolytic synapse formation. Together, these data suggest that, beyond the well-established inhibitory roles of PKA type I, compartmentalized increases in cAMP capable of selectively activating PKA type II in the AKAP350 complex may exert a facilitatory effect on NK-cell cytotoxicity by promoting CIP4 translocation to the IS.

Our *in-situ* studies showed that the non-phosphorylable CIP4 T225A mutant showed a significantly higher association to microtubules that the phosphomimetic CIP4 T225E mutant, which suggests that CIP4 phosphorylation by PKA directly induces CIP4 delocalization from microtubules. Regarding the mechanisms underlying those observations, AlphaFold’s iPTM prediction for theCIP4 dimer phosphorylated at residue T225 result was markedly higher compared to that for the non-phosphorylated dimer. The elevated iPTM value can be interpreted as a more conformationally constrained dimeric conformation, suggesting that T225 phosphorylation alters CIP4 intermolecular architecture in a way that may influence its functional interactions during NK cell activation. CIP4 dimer has been shown to be the structural form that interacts with the plasma membrane and drives membrane curvature (Shimada et al., 2007). In this context, our previous work demonstrated that the T225 phosphomimetic mutant displays increased localization at invadopodia compared to the non-phosphorylatable variant (Tonucci et al., 2019), thus supporting an effect of CIP4T225 phosphorylation on its location in protrusive membrane structures. Whether phosphorylation of CIP4 at T225 directly facilitates its positioning at the IS by stabilizing CIP4 dimer interaction, and thereby contributes to its role in coordinating cytoskeletal organization with IS membrane remodeling will require further investigation.

In summary, this study identifies a signaling mechanism that regulates the subcellular localization of CIP4 during NK-cell activation and provides mechanistic insight into how cytoskeletal coordination is achieved during NK-IS formation. We demonstrate that NK-cell interaction with sensitive targets enhances CIP4 phosphorylation and that disruption of the PKA/AKAP350 pathway prevents CIP4 relocalization to the IS. Together with the differential microtubule association of non-phosphorylatable and phosphomimetic CIP4 T225 mutants, these findings establish phosphorylation at T225 as a key determinant of CIP4 spatial regulation during synapse assembly. By integrating upstream signaling events with the dynamic positioning of a cytoskeletal scaffold protein, our results underscore the relevance of compartmentalized cAMP–PKA signaling in the control of NK response during the establishment of a cytolytic IS.

## ACKNOWLEDGEMENTS AND FUNDING

This work was supported by CONICET Grant PIP2720 and ANPCyT grant PICT2020-02817 (to MCL). The funders had no role in study design, data collection and analysis, decision to publish, or paper preparation. We gratefully acknowledge Dr. Norberto Zwirner for his expert guidance in developing the protocol for ex vivo NK cell purification and genetic manipulation, as well as for generously providing the essential reagents necessary for these procedures.

## DATA AVAILABILITY

The datasets generated during the current study are available from the corresponding author on reasonable request.

---

**Supplementary Figure 1. AlphaFold 3 predicted structure of CIP4 dimer non-phosphorylated at T225**. A) General view of the dimer colored by pLDDT per-atom (yellow < pLDDT < blue). Figures were generated with ChimeraX and colored by alphafold palette (bfactor). B) Predicted aligned error (PAE) graphic for the dimmer. C) Lateral and diagonal views with T225 colored in red.

**Supplementary Figure 2. AlphaFold 3 predicted structure of CIP4 dimer phosphorylated at T225.** A) General view of the dimer colored by pLDDT per-atom (yellow < pLDDT < blue). B) Figures were generated with ChimeraX and colored by alphafold palette (bfactor). Predicted aligned error (PAE) graphic for the dimmer. C) Lateral and diagonal views with phosphorylated T225 colored in red.

**Supplementary Figure 3.** NK–target cell conjugates were formed by incubating NK-YTS and KT86 cells at a 2:1 ratio for 30 min, followed by fixation and staining. CIP4 association with microtubules was quantified as described in the Materials and Methods section. A) Conjugate formation was performed under control conditions or in the presence of H89 (1 μM), as described in Figure 4. B) Conjugates were formed using control or AKAP350KD NK-YTS cells, as described in Figure 5. Dot plots show quantification from three independent experiments of the percentage of CIP4 associated with microtubules in isolated (resting) NK-YTS cells and in NK cells conjugated to target cells (IS) under control conditions or in the presence of H89 (A) or following AKAP350 knockdown (B). Data are presented as mean ± SEM.

