## Supplementary Figures 1 and 2 for "PKA–CIP4 SIGNALING REGULATES CIP4 RELOCATION IN ACTIVATED NATURAL KILLER CELLS"

### Supplementary figure 1

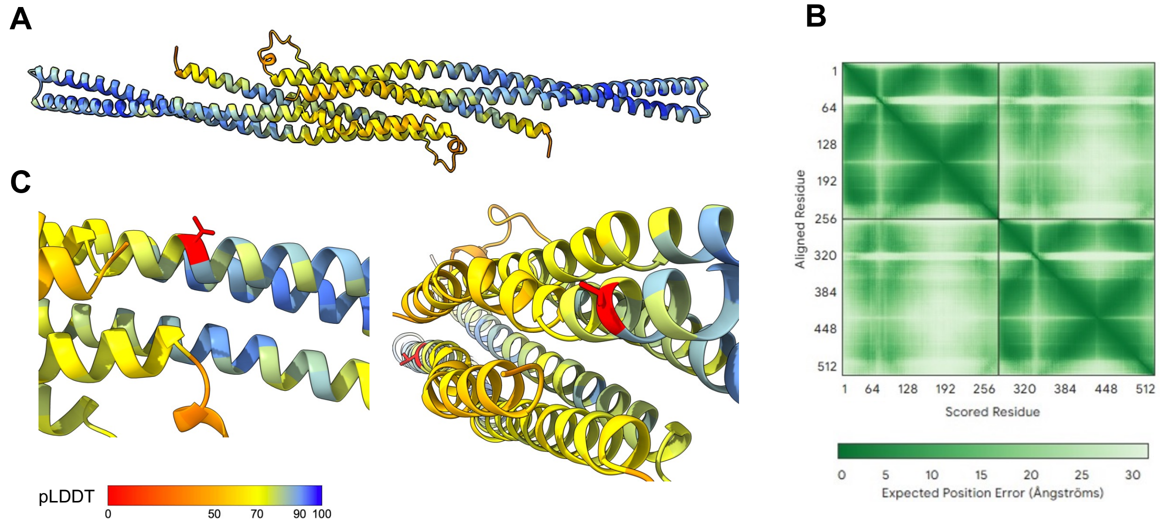


**AlphaFold 3 predicted structure of CIP4 dimmer non phosphorilated at T225**. A) General view of the dimmer colored by pLDDT per-atom (yellow < pLDDT < blue). Figures were generated with ChimeraX and colored by alphafold palette (bfactor). B) Predicted aligned error (PAE) graphic for the dimmer. C) Lateral and diagonal views with T225 colored in red.

### Supplementary figure 2

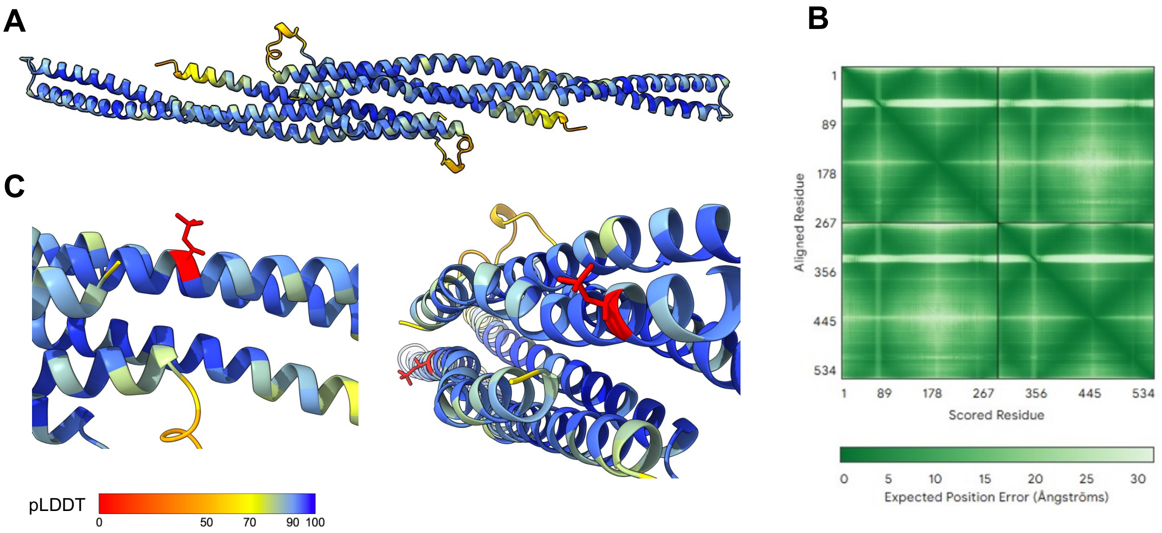


**AlphaFold 3 predicted structure of CIP4 dimmer phosphorilated at T225**. A) General view of the dimmer colored by pLDDT per-atom (yellow < pLDDT < blue). B) Figures were generated with ChimeraX and colored by alphafold palette (bfactor). Predicted aligned error (PAE) graphic for the dimmer. C) Lateral and diagonal views with T225P colored in red.

**Supplementary Figure 3**

A

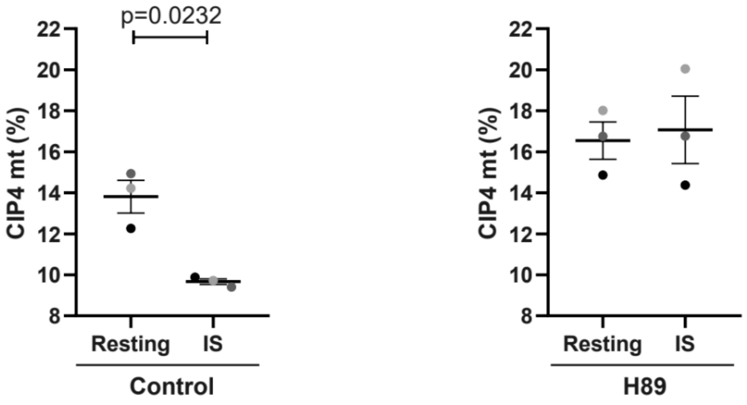


B

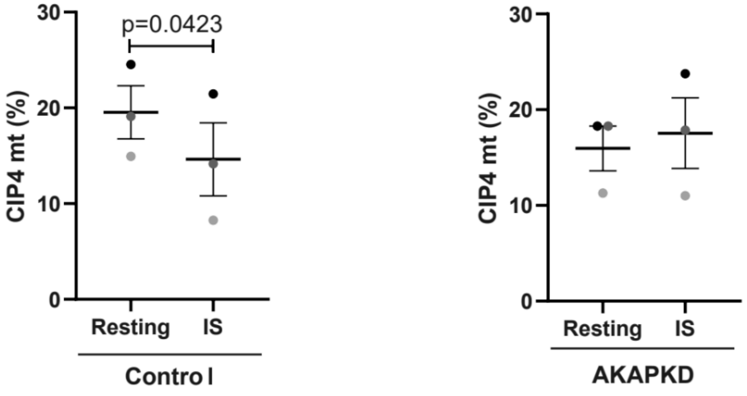


NK–target cell conjugates were formed by incubating NK-YTS and KT86 cells at a 2:1 ratio for 30 min, followed by fixation and staining. CIP4 association with microtubules was quantified as described in the Materials and Methods section. A) Conjugate formation was performed under control conditions or in the presence of H89 (1 µM), as described in Figure 4. B) Conjugates were formed using control or AKAP350KD NK-YTS cells, as described in Figure 5. Dot plots show quantification from three independent experiments of the percentage of CIP4 associated with microtubules in isolated (resting) NK-YTS cells and in NK cells conjugated to target cells (IS) under control conditions or in the presence of H89 (A) or following AKAP350 knockdown (B). Data are presented as mean ± SEM. Statistical significance was determined using a paired t test.
